# Machine Learning to Predict Osteoporotic Fracture Risk from Genotypes

**DOI:** 10.1101/413716

**Authors:** Vincenzo Forgetta, Julyan Keller-Baruch, Marie Forest, Audrey Durand, Sahir Bhatnagar, John Kemp, John A Morris, John A Kanis, Douglas P. Kiel, Eugene V McCloskey, Fernando Rivadeneira, Helena Johannson, Nicholas Harvey, Cyrus Cooper, David M Evans, Joelle Pineau, William D Leslie, Celia MT Greenwood, J Brent Richards

## Abstract

**Background:** Genomics-based prediction could be useful since genome-wide genotyping costs less than many clinical tests. We tested whether machine learning methods could provide a clinically-relevant genomic prediction of quantitative ultrasound speed of sound (SOS)—a risk factor for osteoporotic fracture.

**Methods:** We used 341,449 individuals from UK Biobank with SOS measures to develop genomically-predicted SOS (gSOS) using machine learning algorithms. We selected the optimal algorithm in 5,335 independent individuals and then validated it and its ability to predict incident fracture in an independent test dataset (N = 80,027). Finally, we explored whether genomic pre-screening could complement a UK-based osteoporosis screening strategy, based on the validated tool FRAX.

**Results:** gSOS explained 4.8-fold more variance in SOS than FRAX clinical risk factors (CRF) alone (*r*^*2*^ = 23% vs. 4.8%). A standard deviation decrease in gSOS, adjusting for the CRF-FRAX score was associated with a higher increased odds of incident major osteoporotic fracture (1,491 cases / 78,536 controls, OR = 1.91 [1.70-2.14], P = 10^-28^) than that for measured SOS (OR = 1.60 [1.50-1.69], P = 10^-52^) and femoral neck bone mineral density (147 cases / 4,594 controls, OR = 1.53 [1.27-1.83], P = 10^-6^). Individuals in the bottom decile of the gSOS distribution had a 3.25-fold increased risk of major osteoporotic fracture (P = 10^-18^) compared to the top decile. A gSOS-based FRAX score, identified individuals at high risk for incident major osteoporotic fractures better than the CRF-FRAX score (P = 10^-14^). Introducing a genomic pre-screening step into osteoporosis screening in 4,741 individuals reduced the number of required clinical visits from 2,455 to 1,273 and the number of BMD tests from 1,013 to 473, while only reducing the sensitivity to identify individuals eligible for therapy from 99% to 95%.

**Interpretation:** The use of genotypes in a machine learning algorithm resulted in a clinically-relevant prediction of SOS and fracture, with potential to impact healthcare resource utilization.

**Research in Context:** *Evidence Before this Study:* Genome-wide association studies have identified many loci associated with risk of clinically-relevant fracture risk factors, such as SOS. Yet, it is unclear if such information can be leveraged to identify those at risk for disease outcomes, such as osteoporotic fractures. Most previous attempts to predict disease risk from genotypes have used polygenic risk scores, which may not be optimal for genomic-prediction. Despite these obstacles, genomic-prediction could enable screening programs to be more efficient since most people screened in a population are not determined to have a level of risk that would prompt a change in clinical care. Genomic pre-screening could help identify individuals whose risk of disease is low enough that they are unlikely to benefit from screening.

*Added Value of this Study:* Using a large dataset of 426,811 individuals we trained and tested a machine learning algorithm to genomically-predict SOS. This metric, gSOS, had performance characteristics for predicting fracture risk that were similar to measured SOS and femoral neck BMD. Implementing a gSOS-based pre-screening step into the UK-based osteoporosis treatment guidelines reduced the number of individuals who would require screening clinical visits and skeletal testing by approximately 50%, while having little impact on the sensitivity to identify individuals at high risk for osteoporotic fracture.

*Implications of all of the Available Evidence:* Clinically-relevant genomic prediction of heritable traits is feasible using the machine learning algorithm presented here in large sample sizes. Genome-wide genotyping is now less expensive than many clinical tests, needs to be performed once over a lifetime and could risk stratify for multiple heritable traits and diseases years prior to disease onset, providing an opportunity for prevention. The implementation of such algorithms could improve screening efficiency, yet their cost-effectiveness will need to be ascertained in subsequent analyses.

## Introduction

Genomic prediction of complex traits and diseases could improve the efficiency of health care systems if it can accurately identify those at risk for disease. This potential arises because genotypes are stable over the lifetime so that high risk individuals might be identified early in life, providing an opportunity for prevention years before disease onset. Further, genotypes need to be measured only once over a lifetime and could provide risk prediction for many heritable diseases. Finally, since the cost of genome-wide genotype is now reasonably affordable (approximately $40 in a research context) and falling faster than Moore’s law,^1^ genotype-based prediction has become less expensive than some clinical tests. For these reasons genome-wide genotyping has already been undertaken in several health care contexts.^2,3^ Yet despite this promise, it remains unclear if genomics can render clinically-relevant predictions and perhaps more importantly, how such information could be used to improve care.

The use of genetic prediction can be explored using osteoporosis as an example, since it is among the most common diseases and is diagnosed primarily through bone density measures, which are highly heritable (50-85%) and highly polygenic.^4–7^ Osteoporosis is defined as a reduction in bone mass and microarchitectural integrity resulting in an increased risk of fracture.^8^ Many guideline-based screening strategies^9–12^ for prevention of osteoporosis-related fractures incorporate the Fracture Risk Assessment Tool (FRAX),^13,14^ a validated method to risk stratify individuals for treatment. Further, a FRAX-based screening program has also been shown to reduce hip fracture incidence.^15^ However, population-based screening programs identify a small proportion of the screened population that is eligible for lifestyle or treatment interventions and require relatively expensive BMD testing in a large proportion of the population who will not ultimately receive treatment. Thus, current osteoporosis care models provide the opportunity to explore how genomic prediction may provide clinical utility.

At present, there is little evidence that genotype-based prediction algorithms provide clinical utility for common disease. Since the variance explained by genotypes in complex traits and diseases by genotypes has been typically low, the resulting performance of genetic prediction has not rendered important improvements over clinical risk factors.^16,17^ One way to improve the variance explained by genotypes is through the use of larger sample sizes, applying genome-wide association study (GWAS) polygenic risk scores,^18–21^ since for many complex traits and diseases the effect sizes of any one associated genetic risk factor is small, but taken together, the variance explained can be substantial.^22^ Machine-learning algorithms may also be applied to larger sample sizes to improve prediction, where these methods learn from training data and then assess performance of the algorithm in independent test datasets.^23^

In this study, we trained a machine learning algorithm and separate polygenic risk scores on a large cohort of 341,449 individuals of British ancestry from UK Biobank with genome-wide genotypes to predict bone quantitative ultrasound-derived speed of sound (SOS) a measure that predicts osteoporotic fracture partly independently of DXA-derived bone mineral density (BMD) and clinical risk factors.^24^ We then selected the optimal prediction model in 5,335 independent individuals and finally tested the clinically-relevant performance characteristics of genomically-predicted SOS (gSOS) for fracture prediction in 80,027 independent individuals. Finally, we explored how gSOS could be incorporated into FRAX-based screening strategies in order to identify a proportion of the population not needing screening and/or BMD testing.

## Methods

### Overall Study Design

This study included three phases (**Figure 1**). The first was to use machine learning methods^25^ and polygenic risk scores to predict SOS in the training dataset, selecting the top-performing model in the model selection dataset, which we refer to as gSOS. The second phase used the independent test dataset to assess the ability of gSOS to predict fracture and the third was to explore how gSOS may be used to improve screening programs.

**Figure 1.**
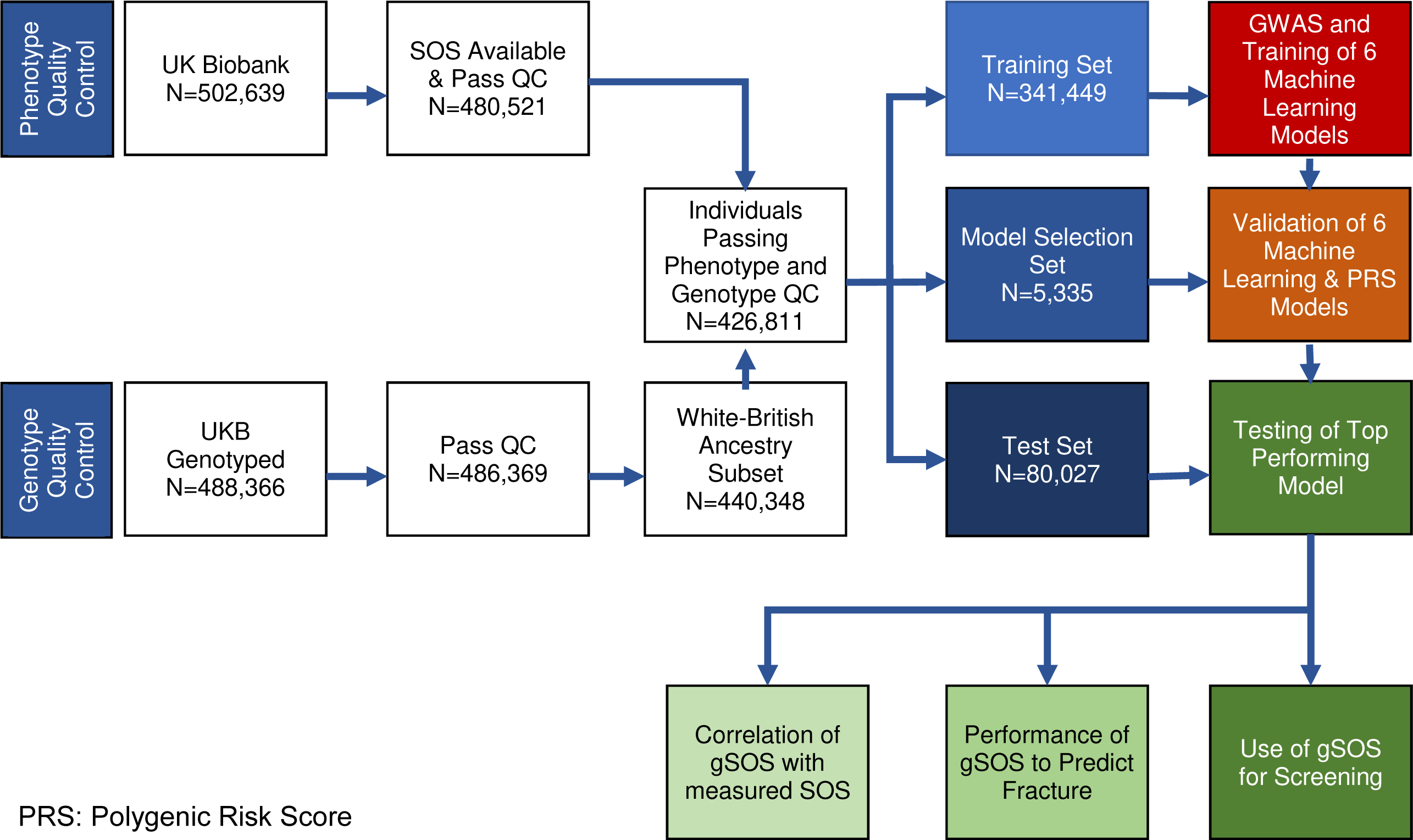
Overall Study Design.

### Cohort

UK Biobank is a large-scale health resource that follows 502,628 volunteer participants in the United Kingdom.^26^ Participants within the UK Biobank have been genome-wide genotyped using Affymetrix arrays,^27^ followed by genotype imputation to the Haplotype Reference Consortium.^28^ UK Biobank has ethical approval from the Northwest Multi-centre Research Ethics Committee, and informed consent was obtained from all participants.

### SOS and BMD Measurement (Details in Supplement)

Details of SOS measurement are available from UK Biobank (<u>goo.gl/A5DTdP)</u>. All analyses used SOS standardized to a mean of zero and standard deviation of one. We chose to model SOS, since it was available for more individuals than BMD, has a gradient of risk for incident major osteoporotic fractures similar to BMD and a recent meta-analysis described its effect on fracture risk, allowing for the development of a SOS-based FRAX model.^24^ Dual energy x-ray absorptiometry BMD was measured at the femoral neck (and is referred to hereafter as “BMD”) in a subset of the population (N = 4,741), that was reserved for the test dataset in order to assess osteoporosis screening strategies.

### Development of machine learning model to predict SOS (Supplement and Figure 1)

#### I. Training, Model Selection and Test datasets

To develop and test our prediction models, we followed best practices in machine learning to ensure unbiased estimates of model performance. Participants in UK Biobank with White British ancestry, measured SOS, and genotyping information (N= 426,811) were randomly assigned to either a training dataset (80% of participants, restricted to those without BMD), a model selection dataset (1.25%, also without BMD), or the test dataset (18.75% of participants including all with BMD) (**Figure 1** & **Table 1**).

**Table 1:**
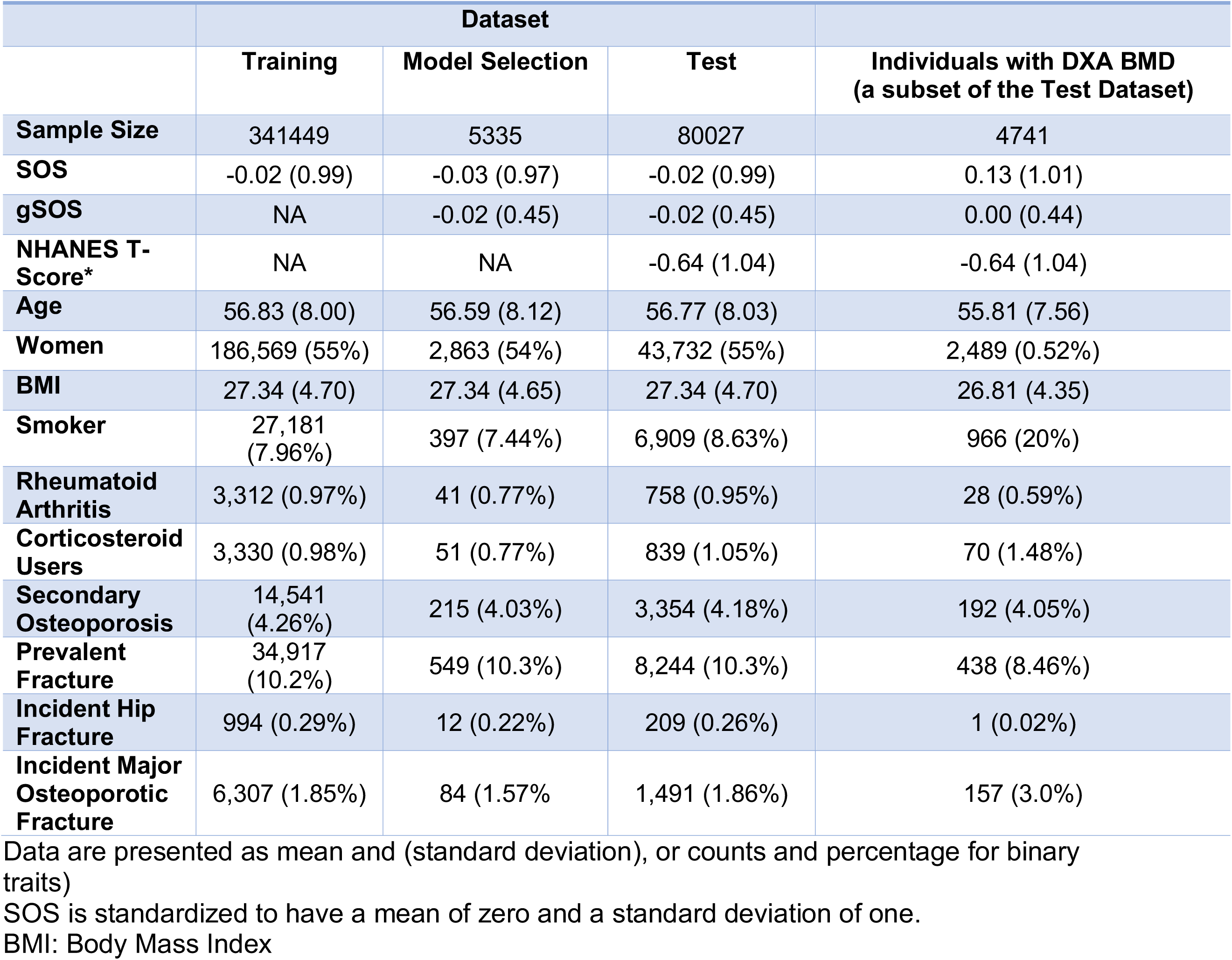
Characteristics of Participants by Dataset.

|  | Dataset |  |  | Individuals with DXA BMD<br>(a subset of the Test Dataset) |
| --- | --- | --- | --- | --- |
|  | Training | Model Selection | Test |  |
| **Sample Size** | 341449 | 5335 | 80027 | 4741 |
| **SOS** | -0.02 (0.99) | -0.03 (0.97) | -0.02 (0.99) | 0.13 (1.01) |
| **gSOS** | NA | -0.02 (0.45) | -0.02 (0.45) | 0.00 (0.44) |
| **NHANES T-Score\*** | NA | NA | -0.64 (1.04) | -0.64 (1.04) |
| **Age** | 56.83 (8.00) | 56.59 (8.12) | 56.77 (8.03) | 55.81 (7.56) |
| **Women** | 186,569 (55%) | 2,863 (54%) | 43,732 (55%) | 2,489 (0.52%) |
| **BMI** | 27.34 (4.70) | 27.34 (4.65) | 27.34 (4.70) | 26.81 (4.35) |
| **Smoker** | 27,181<br>(7.96%) | 397 (7.44%) | 6,909 (8.63%) | 966 (20%) |
| **Rheumatoid Arthritis** | 3,312 (0.97%) | 41 (0.77%) | 758 (0.95%) | 28 (0.59%) |
| **Corticosteroid Users** | 3,330 (0.98%) | 51 (0.77%) | 839 (1.05%) | 70 (1.48%) |
| **Secondary Osteoporosis** | 14,541<br>(4.26%) | 215 (4.03%) | 3,354 (4.18%) | 192 (4.05%) |
| **Prevalent Fracture** | 34,917<br>(10.2%) | 549 (10.3%) | 8,244 (10.3%) | 438 (8.46%) |
| **Incident Hip Fracture** | 994 (0.29%) | 12 (0.22%) | 209 (0.26%) | 1 (0.02%) |
| **Incident Major Osteoporotic Fracture** | 6,307 (1.85%) | 84 (1.57%) | 1,491 (1.86%) | 157 (3.0%) |

Data are presented as mean and (standard deviation), or counts and percentage for binary traits)
SOS is standardized to have a mean of zero and a standard deviation of one.
BMI: Body Mass Index

#### II. GWAS

Using the training dataset (N=341,449 individuals with White British ancestry), we tested the additive allelic effects of each of the 13M SNPs passing QC, separately, on SOS using a series of linear mixed-model^29^, adjusting for age, sex, assessment centre, genotyping array, and the first 20 principal components derived from the analysis of White British ancestry. Linkage disequilibrium-independent associations where obtained using PLINK by clumping SNPs in linkage equilibrium at a r^2^ > 0.05 and selecting a single most significant SNP from within each clumped set.

#### III. Machine learning models

We fit 6 regularized linear machine learning models^25^ based on L1-penalized least absolute shrinkage and selection operator (LASSO) regression^25^ to the training dataset to predict SOS using only SNPs with P-values smaller than a chosen set of thresholds (**Table S2**). The model selection dataset was then used to optimize model performance by choosing the best regularization parameter (λ) which controls the number of SNPs contributing to the prediction. Finally, the estimated model for the SNP-set giving the smallest root mean square error in the model selection dataset was taken forward for testing in the test dataset. Hence, we ensured that the performance of only one final model from the machine learning strategy was evaluated in the test dataset. We refer to this final predictor as “gSOS”, in which SOS is predicted only from genotypes. All evaluations of the role of gSOS on screening tests were performed in the test dataset.

#### IV Polygenic Risk Scores

Traditional polygenic risk scores^18^ were generated summarizing different sets of SNPs, selected by P-value threshold, as described in the **Supplement**.

### Measurement of FRAX risk factors (Details in Supplement)

The FRAX risk score for fracture can be generated with or without BMD, referred to BMD-FRAX and clinical risk factor CRF-FRAX, respectively. FRAX CRFs were assessed at the baseline visit and included age, sex, body mass index (BMI), previous fracture, smoking, corticosteroid use, rheumatoid arthritis and secondary causes of osteoporosis. More than three daily units of alcohol and parental history of hip fracture variables were not available and were set to “no” for the entire population. Age was recorded at baseline visit, and sex was self-reported and verified by genotype. Individuals with discordant sex between self-report and genotype were excluded.

### Fracture Definitions (Details in Supplement)

Site-specific major osteoporotic fractures^30^ were defined using questionnaire data from UK Biobank and ICD10 codes for hospital admissions. For ICD10 codes (**Table S1**), only primary diagnoses were considered, since the date of secondary diagnoses was not consistently available. The date of ICD-10 identified fractures was determined from administrative data and if dated after the baseline visit was labelled as incident. Prevalent fractures were identified as those reported by questionnaire or from ICD10 codes that occurred prior to the baseline visit. Trauma type was not included since mechanism of trauma is variably captured by ICD codes and even high trauma fractures are predictive of future low trauma fractures and are associated with low BMD.^31,32^

### Generation of FRAX scores

Four FRAX scores were generated for each person in the test dataset: 1) CRF-FRAX; 2) CRF plus measured-SOS FRAX (termed SOS-FRAX); 3) CRF plus gSOS FRAX (termed gSOS-FRAX) and 4) CRF plus BMD FRAX (termed BMD-FRAX). CRF-FRAX and BMD-FRAX were generated using the FRAX algorithm.^33^ To generate the SOS-FRAX probabilities, we took the CRF-FRAX score (as pre-test odds) and updated this using measured SOS or gSOS (gradient of risk per standard deviation decrease of 1.42 for major osteoporotic fractures and 1.80 for hip fracture from the largest meta-analysis to date which did not include UK Biobank).^24^ We used this method since a SOS-FRAX score is currently not available.

### Non-White British, Population Stratification and Cryptic Relatedness

The training, model selection and test datasets included only White British participants to reduce population stratification. The GWAS was also controlled for the top 20 principle components of ancestry to reduce effects of cryptic relatedness. To test the generalizability of gSOS and assess for potential effects of population stratification and cryptic relatedness we tested gSOS on fracture outcomes in a smaller cohort of non-White British participants (defined in the **Supplement**).

### Testing of Prediction Performance in the Test Dataset

Prediction performance was first tested by assessing the correlation between gSOS and measured SOS in the test dataset. Next, the correlations between gSOS and FRAX CRFs were assessed. gSOS was tested for association with risk of incident major osteoporotic fracture and hip fracture, comparing this to measured SOS and BMD measured at the femoral neck using univariate and multivariable logistic regression, including CRF-FRAX as a covariate. The association of each of the four FRAX scores with risk of incident major osteoporotic fracture was also tested in the test dataset by first log-transforming the FRAX scores to correct for a skewed distribution and then using logistic regression. The area under the receiver operator curve was calculated for each of the FRAX scores, and these were compared to the CRF-FRAX. The same procedure was then completed for incident hip fracture.

Since health care systems may want to use genomics-based predictors to identify an extreme of the population at risk of disease, we next divided the distribution of gSOS into percentiles and contrasted the bottom decile with the top decile to describe the associated increased risk of fracture. Since 10% of any health care system is a large number of people, we then explored odds of fracture for those in the lowest 25^th^ quantile (lowest 4% of the distribution), compared to the top 25^th^ quantile.

### Genomic Pre-Screening

Health care providers may also want to use genomics-based predictors to better stratify individuals for inclusion in screening programs, assuming that genome-wide genotyping has been already undertaken. We therefore explored if gSOS could be used to identify individuals who would be unlikely to benefit from screening for osteoporosis, because their risk of fracture would be low. To do so, we implemented the UK National Osteoporosis Guideline Group (NOGG) approach within the test dataset.^10^ The NOGG Guidelines aim to identify individuals at risk for fracture in a cost-efficient manner by reserving clinical visits and BMD testing for those at increased risk.

We first evaluated the effectiveness of NOGG Guidelines to appropriately risk stratify individuals for therapy, comparing this to a gold standard where all individuals had BMD testing and were risk stratified by BMD-FRAX. We chose NOGG over other national guidelines since a NOGG-like approach was recently demonstrated in a randomized controlled trial to be potentially effective in reducing hip fractures.^15^ Further, the NOGG Guidelines recommend less BMD testing than those of the US National Osteoporosis Foundation^11^ and Osteoporosis Canada.^34^ Consequently an improvement in BMD testing efficiency over NOGG would suggest improvements in other guideline settings. NOGG-guidelines utilise 10-year absolute risks of fracture using FRAX and suggest treatment or reassurance based on thresholds of risk, which are age dependent and consider competing risks. These 10-year absolute risks were calculated for all individuals for major osteoporotic fracture and participants were risk stratified as per NOGG Guidelines (**Figure S4**), which aims to reduce the number of BMD tests by restricting such tests to those at moderate risk. We then tested the sensitivity of NOGG guidance to identify individuals at high risk, comparing NOGG guidance to a BMD-FRAX gold-standard. Since BMD is required to calculate BMD-FRAX, the genomic pre-screening was evaluated only in individuals from the test dataset with BMD measures (N = 4,741).

In order to implement genomic pre-screening, we designated all individuals over a defined threshold of gSOS to be reassured, and not to undergo a clinical visit for CRF-based FRAX testing, or BMD testing (**Figure S5**). We chose the threshold of gSOS which maximally reduced clinical visits and BMD testing, while minimizing the loss of sensitivity to identify individuals who would be recommended for treatment by the gold standard. The performance of pre-screening was then assessed by counting the number of individuals who would require a clinical visit and BMD testing and calculating its sensitivity to correctly assign participants to reassurance or treatment.

## Results

### Cohort Characteristics

**Table 1** describes the FRAX risk factors for the training, model selection and test data sets. There was no clinically-relevant difference in any of the osteoporosis-related risk factors, as expected, since these sets were generated randomly. As planned, all individuals with BMD measures were included in the test dataset. There were little differences in demographics or CRFs among individuals with BMD measured, except for a higher prevalence of smoking and incident major osteoporotic fractures.

### GWAS

After quality control, 13,958,249 SNPs were included in the GWAS. The GWAS in the training set identified 1,404 independent (*r*^*2*^ ≤ 0.05) genome-wide significant loci at a P-value threshold of < 5 × 10^-8^ **Figure S1** shows the Manhattan and QQ plots for this GWAS.

### Variance Explained in SOS in the Model Selection Set

Age, sex and BMI explained 4.0% of the variance in SOS. Adding the remaining FRAX clinical risk factors increased the variance explained to 4.8%. Polygenic risk scores across different P-value thresholds explained at most 18.5% (95% confidence interval [CI]: 16.6-20.4%) of the variance in SOS (**Table S2** and **Figure 2**). The machine-learning algorithm improved the variance explained in SOS to a maximum of 25.0% (23.0-27.0%) (**Table S2** and **Figure 2**). All six SNP-sets using the machine learning algorithm outperformed polygenic risk scores substantially. **Figure S2** provides detailed information on the optimal algorithm tuning parameters.

**Figure 2.**
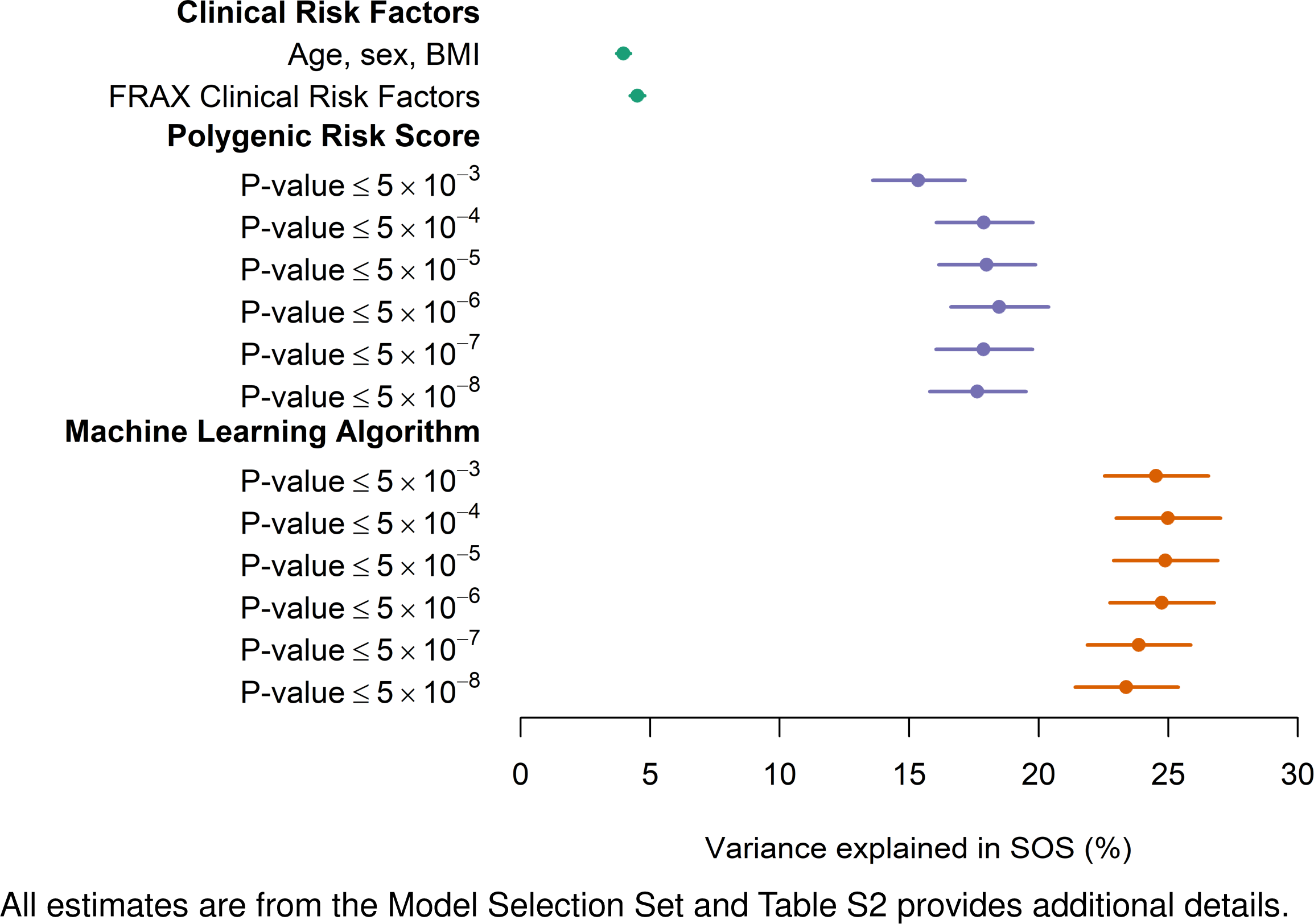
Variance Explained in SOS by Clinical Risk Factors, Polygenic Risk Score and Machine Learning Algorithm in the Model Selection Set.

### Variance Explained in SOS in the Test Set

The top model from the model selection set was then tested for its correlation with SOS in the test set. This model (machine learning algorithm set at P-value ≤ 10^-4^, including 21,717 activated SNPs, which are those SNPs retained by the machine learning algorithm) explained 23.2% (95% CI: 22.7-23.7%) of the variance in measured SOS (**Table S2** and **Figure 2**). We then designated this model as “gSOS”, which was subsequently assessed for performance characteristics in the test dataset. **Figure S3** demonstrates that, as expected, the estimated effects of the activated SNPs from the machine learning algorithm were attenuated, when compared to the effects estimated from the GWAS

### gSOS Correlation with other Clinical Risk Factors and BMD

Since gSOS was derived only from genotypes it should be independent of factors that are not in the causal pathway between SNPs and SOS. We found that while measured SOS was correlated with all measured FRAX clinical risk factors, gSOS was not (**Table 2**). These findings suggest that if gSOS is associated with risk of fracture it would be also independent of these other clinical risk factors.

**Table 2.**
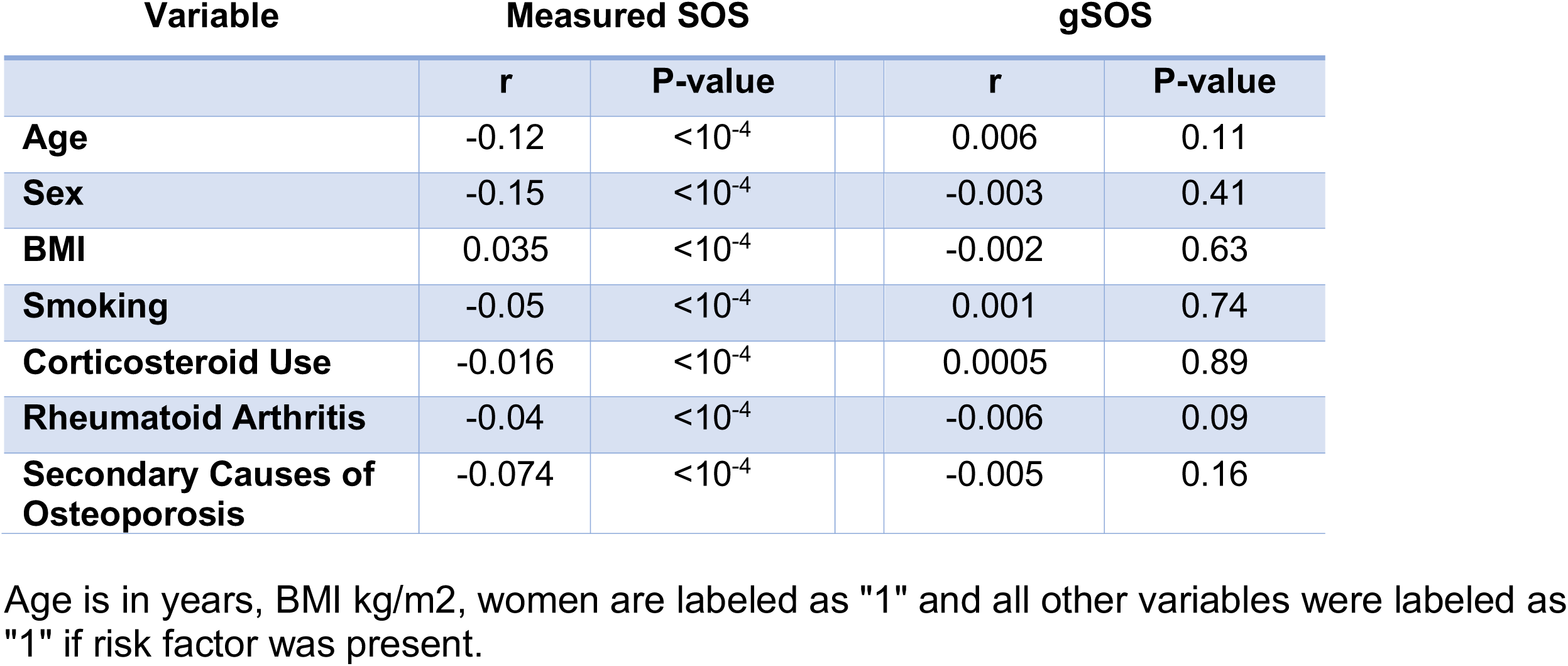
Correlation of gSOS with FRAX Clinical Risk Factors.

| Variable | Measured SOS |  | gSOS |  |
| --- | --- | --- | --- | --- |
|  | r | P-value | r | P-value |
| **Age** | -0.12 | $<10^{-4}$ | 0.006 | 0.11 |
| **Sex** | -0.15 | $<10^{-4}$ | -0.003 | 0.41 |
| **BMI** | 0.035 | $<10^{-4}$ | -0.002 | 0.63 |
| **Smoking** | -0.05 | $<10^{-4}$ | 0.001 | 0.74 |
| **Corticosteroid Use** | -0.016 | $<10^{-4}$ | 0.0005 | 0.89 |
| **Rheumatoid Arthritis** | -0.04 | $<10^{-4}$ | -0.006 | 0.09 |
| **Secondary Causes of Osteoporosis** | -0.074 | $<10^{-4}$ | -0.005 | 0.16 |

Age is in years, BMI kg/m<sup>2</sup>, women are labeled as "1" and all other variables were labeled as "1" if risk factor was present.

### Prediction of Incident Major Osteoporotic and Hip Fracture

There were 1,491 incident major osteoporotic fracture cases and 78,536 incident fracture-free controls in the test dataset (**Figure 3**). We also identified 209 hip fracture cases and 79,818 incident fracture-free controls in the same dataset. In univariate models, decreased SOS, gSOS and BMD were all strongly associated with increased odds of incident fracture, however, among these predictors, gSOS had the highest risk per standard deviation decrease. Since the predictive value of these skeletal measures may be reduced after accounting for correlated FRAX CRFs, we tested the association of SOS, gSOS and BMD with incident major osteoporotic and hip fractures, adjusting for CRF-FRAX. We found that association of fracture with SOS and BMD were more attenuated by inclusion CRF-FRAX (**Figure 3**), likely because gSOS is not associated with FRAX CRFs, while SOS and BMD are. Associations with incident hip fractures showed similar effect sizes, but with wider confidence intervals. There were too few incident hip fractures amongst the 4,741 participants with BMD measures to report meaningful findings.

**Figure 3.**
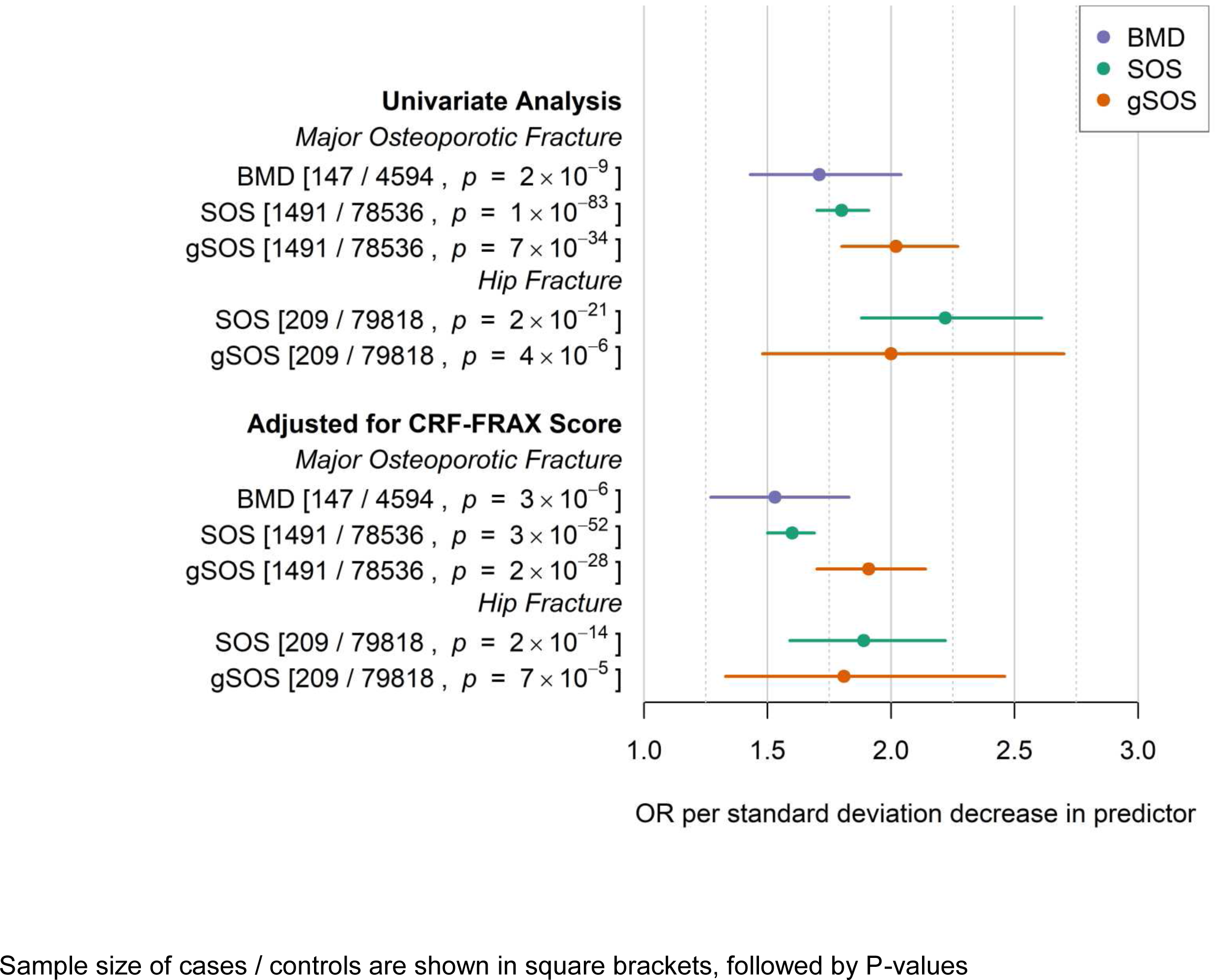
Performance of SOS, gSOS and BMD for the Prediction of Major Osteoporotic and Hip Fractures in the Test Dataset.

### Incorporating gSOS into FRAX Probability of Fracture

We next tested the performance of the four FRAX probabilities on risk of incident major osteoporotic fracture. We found that a standard deviation increase in each FRAX score on the log scale was strongly associated with incident major osteoporotic fracture (**Table 3**). The magnitude of association was lower when using CRFs alone and increased when introducing any of the three skeletal measures. The area under the receiver operator curves improved significantly, with the incorporation of skeletal measures when compared to CRFs alone, as has been shown previously.^33^ **Table 3** shows that the area under the receiver operator curve for gSOS-FRAX was equivalent to that for BMD-FRAX but was lower than SOS-FRAX.

**Table 3.** Prediction of Incident Fractures by FRAX Scores.

| Major Osteoporotic Fracture |  |  |  | Hip Fractures |  |  |  |
| --- | --- | --- | --- | --- | --- | --- | --- |
| Predictor | Odds per 1 log increase in predictor (95% CI) | AUC (95% CI) | Comparison of AUC with gSOS AUC<sup>a</sup> |  | Odds per 1 log increase in predictor (95% CI) | AUC (95% CI) | Comparison of AUC with gSOS AUC<sup>a</sup> |
| CRF-FRAX | 1.68<br>(1.60-1.76) | 65%<br>(64%-66%) | gSOS improves AUC<br>(P = 3x10<sup>-14</sup>) |  | 2.95<br>(2.54-3.42) | 77%<br>(74%-80%) | gSOS improves AUC<br>(P = 4x10<sup>-3</sup>) |
| BMD-FRAX | 1.69<br>(1.47-1.96) | 66%<br>(62%-71%) | gSOS equivalent to BMD<br>(P = 0.47) |  | N/A | N/A | N/A |
| SOS-FRAX | 1.91<br>(1.82-2.01) | 68%<br>(67%-70%) | SOS improves AUC<br>(P = 7x10<sup>-9</sup>) |  | 2.67<br>(2.36-3.02) | 79%<br>(76%-82%) | SOS equivalent to gSOS<br>(P = 0.09) |
| gSOS-FRAX | 1.77<br>(1.69-1.86) | 67%<br>(65%-68%) | - |  | 2.94<br>(2.55-3.39) | 78%<br>(75%-81%) | - |

N/A: Too few cases of hip fractures amongst 4741 individuals with BMD measures
CRF-FRAX: Clinical Risk Factors FRAX Score
BMD-FRAX: Clinical Risk Factors plus BMD FRAX Score
SOS-FRAX: Clinical Risk Factors plus SOS FRAX Score
gSOS-FRAX: Clinical Risk Factors plus gSOS FRAX Score
AUC: Area under the receiver operator curve
a) Comparison with gSOS compares the AUC for gSOS with each of the other predictors

We next generated the FRAX probabilities using hip fracture as an outcome. **Table 3** shows that each of the models was strongly associated with odds of incident hip fracture and both SOS-FRAX and gSOS-FRAX improved the area under the curve when compared to CRF-FRAX. The area under the curve for SOS-FRAX and gSOS-FRAX was equivalent.

### Extremes of gSOS

Comparing individuals in the bottom decile of the gSOS distribution to those in the top decile we found that those in the bottom were at a 3.25-fold increased risk of major osteoporotic fracture (95% CI: 2.50-4.25, P = 10^-18^). Since a decile represents a large absolute number of individuals from any health care system we next tested the risk associated with being in the bottom 25^th^ of the gSOS distribution, compared to those in the top 25^th^ of the distribution and found a ∼5-fold increased risk of incident major osteoporotic fracture (OR = 4.94 [3.07-7.91], P = 10^-11^).

### Non-White British

To assess for generalizability of gSOS and possible effects of population stratification and cryptic relatedness we tested its performance in non-White British participants. There were 494 incident major osteoporotic fracture cases and 42,902 controls amongst non-White British participants. There were 69 incident hip fracture cases and 43,327 controls. The results for non-White British participants were very similar to those amongst White British participants (**Table S3**). gSOS was associated with a higher odds of major osteoporotic and hip fracture, both with and without FRAX CRFs. The area under the receiver operator curve was improved using gSOS-FRAX compared to CRF-FRAX for major osteoporotic and hip fracture (P = 1×10^-4^ and P = 4×10^-4^, respectively). Quantiles of gSOS also demonstrated a high odds of fracture (bottom 10th quantile OR = 3.71 [95% CI: 2.33-5.89, P = 1×10^-8^]; bottom 25^th^ quantile OR = 8.17 [95% CI: 3.22-20.75, P = 5×10^-6^]).

### Genomic Pre-Screening

**Table 4** demonstrates the implementation of current NOGG Guidelines for screening amongst all 4,741 UK Biobank participants with BMD testing. **Figure S4** shows what would be the clinical outcomes (i.e. reassurance or treatment recommendation) based on current NOGG Guidelines. The sensitivity for correct treatment allocation, compared to undertaking BMD-FRAX on all individuals as the gold-standard, was 99.1% and the specificity was 99.6%. To achieve this performance, 2,455 of the 4,741 individuals required a clinical visit while 1,013 required a BMD test (**Figure S4** and **Table 4 and Table S4**).

**Table 4.** Effect of gSOS Pre-Screening on NOGG Screening Tests and Outcomes.

|  |  | Screening:<br>Standard of Care | Screening After<br>Introduction of<br>gSOS Pre-Screen |
| --- | --- | --- | --- |
| *Number of*<br>*Participants* | **Number of**<br>**Participants with**<br>**Sufficient Data**<br>**Available to Test**<br>**Efficacy of Screening** | 4741 |  |
|  | **Number of**<br>**Participants Eligible**<br>**for Screening** | 2455 |  |
| *Clinical*<br>*Visits, BMD*<br>*Tests and*<br>*Outcomes* | **Number of**<br>**Participants**<br>**Requiring a Clinical**<br>**Visit** | 2455 | 1273 |
|  | **Number of**<br>**Participants**<br>**Requiring a BMD Test** | 1013 | 473 |
|  | **Number of**<br>**Participants**<br>**Reassured After**<br>**Screening** | 2223 | 2238 |
|  | **Number of**<br>**Participants**<br>**Recommended for**<br>**Treatment** | 232 | 217 |
| *Performance* | **Sensitivity For**<br>**Correct Treatment**<br>**Allocation<sup>a</sup>** | 99.1% | 94.7% |
|  | **Specificity For**<br>**Correct Treatment**<br>**Allocation** | 99.6% | 99.9% |
|  | **Positive Predictive**<br>**Value** | 96.5% | 98.6% |
|  | **Negative Predictive**<br>**Value** | 99.9% | 99.5% |

Table S4 provides the outcomes of each participant

We then tested whether pre-screening with gSOS could improve efficiency of these guidelines, by reassuring everyone over a certain gSOS threshold. Reassuring all NOGG-eligible participants at a high gSOS threshold would result in little improvement in efficiency.

Alternatively reassuring all participants above too low a gSOS threshold would result in a large number of participants being reassured who would actually be recommended for treatment if they had had their BMD-FRAX measured. Thus, we selected the threshold of gSOS that would minimize the number of BMD tests not changing care, but also not increase the number of participants who are eligible for therapy but would be reassured (**Figure S5**). **Figure S6** and **Table 4** demonstrate that gSOS pre-screening would reduce the number of participants requiring a clinical visit from 2,455 to 1,273 and decrease the number of BMD tests from 1,013 to 473. Using gSOS-based pre-screening, the sensitivity for correct treatment allocation decreased from 99.1% to 94.7% (**Table 4**), while the specificity, positive predictive value and negative predictive value were 99.9%, 98.6% and 99.5%, resptively.

## Discussion

Here we have developed and validated a machine learning algorithm to predict a skeletal measure of fracture risk (gSOS) from genotypes alone. gSOS has performance characteristics for fracture risk that are similar to measured SOS and femoral neck BMD. The machine learning algorithms used here provide improved prediction over traditionally-used polygenic risk scores. While the exact use of such prediction algorithms in clinical care will require further investigation, we have explored several potential scenarios

First, in multivariate models, gSOS was more strongly associated with major osteoporotic fracture than was SOS or BMD. For fracture prediction, gSOS outperformed FRAX CRFs alone. The lower tail of the gSOS distribution was at a substantially increased risk of incident fracture, providing the opportunity to identify individuals at elevated risk. gSOS pre-screening was able to identify a large proportion of the population for whom osteoporosis screening programs would not result in changes in clinical care. These findings suggest that genomic prediction can render clinically-relevant information that could be used to improve health care delivery systems. We suggest that such advances will not be limited to osteoporosis, but rather, given large enough sample sizes and use of machine learning algorithms, genomic prediction will become clinically-relevant for several heritable traits and diseases.

Already at least seven large health care delivery systems have invested in genome-wide genotyping of a large proportion of their population, upon whom electronic health record (EHR) data are available.^2,3^ Since the costs associated with genome-wide genotyping have now dropped below those of many routine clinical tests, the use of genomic prediction of clinically relevant traits and diseases will likely encourage other health care systems to make similar investments—provided genomic predictors can be shown to be clinically relevant and cost efficient.

The fracture prediction performance of gSOS is higher than expected given the amount of variance explained in measured SOS. This is likely for at least three reasons. First, gSOS is not subject to measurement error inherent in SOS; second it is independent of other FRAX clinical risk factors and last, genotypes are measured with very high precision.^35^ It is not currently clear which machine learning methods are best suited to prediction. Nonetheless, the observed association between genotypes and continuous traits has largely been linear, making multivariate regression-based methods, like LASSO, attractive.

Screening programs for osteoporosis are expensive, with estimates of approximately $50,000 USD per quality adjusted life year gained,^36^ but are less expensive using NOGG guidance.^37^ Current guidelines suggest screening for a large proportion of the population,^9–11^ yet most patients are not identified to have a clinically-actionable level of risk.^15,38^ This provides an opportunity for genetically-derived measures of risk to increase cost-efficiencies, particularly in health care systems where investments have been made in genome-wide genotyping. While the health-economic realities of such prediction are unclear at present, we have provided one example of how gSOS could be used to improve the efficiency of screening based on the NOGG Guidelines. We note that the efficiencies of gSOS pre-screening are likely magnified in the context of UK Biobank, where most subjects are generally healthy, yet those receiving BMD tests were more likely to be smokers and have had an incident fracture.

Previous attempts to predict osteoporosis from genomic data did not substantially increase discrimination, when compared to standard clinical measures alone, likely because the GWAS that underpinned these attempts were derived from 32,961 individuals and explained only 5.8% of the variance in BMD.^17,39^ The improvement in variance explained in in this study was likely due to the increase in sample size afforded by UK Biobank as well as the ability of the machine learning algorithm to estimate the joint contributions of all SNPs rather than their independent effects. Other attempts to predict BMD have been based on several dozen genome-wide significant SNPs,^17^ whereas here we have used a machine learning algorithm, which has enabled the consideration of approximately up to 642,127 SNPs. Recently, a machine-learning algorithm was used to predict estimated BMD, but from a GWAS sample size that was one third of that used here.^40^ An additional prediction algorithm using incorrect definitions of osteoporosis was able to explain only 17% of the variance in estimated BMD and was able to predict fracture only marginally better than chance alone.^41^ Therefore the data presented here represent a clinically-relevant improvement in genomic prediction.

The generalizability of genomic prediction can be influenced by population stratification and cryptic-relatedness. Encouragingly, gSOS, despite being developed in the White British subset of UK Biobank, performed similarly in the non-White British subset, who have a different ancestry than the White British subset.

This study has important limitations. While gSOS does risk stratify for fracture, it must be emphasized that this is a probability and is not deterministic—such probabilities have similar characteristics to other risk factors for common disease. BMD would have been a more natural skeletal measure of fracture risk, since this is more often used in the clinic. We anticipate that the same methods here will be useful for BMD genomic prediction, once appropriately large sample sizes are available and emphasize that the predictive performance of gSOS is similar to that for measured BMD. While nearly all FRAX CRFs were available for study, parental hip fracture history was not. It is possible that the clinical relevance of gSOS may be attenuated after accounting for this heritable risk factor, and this will need to be empirically tested once such data are available. gSOS-FRAX could, in the future be optimized to include competing risks and interaction effects, as has been done for BMD-FRAX. Our results have not been validated in an external sample but have been internally validated in 80,027 individuals in UK Biobank. Like participants in most cohort studies, UK Biobank participants are, on average, healthier than the general population.^42^ Thus, external validation in a truly population-based study may provide helpful estimates of predictive abilities of gSOS.

In summary, we have developed and validated a machine-learning algorithm to predict SOS from genotypes alone, which when predicting fracture, performs similarly to measured SOS and BMD. The methods and results provided here may help to guide genomic prediction programs for other clinically-relevant risk factors and diseases, providing new opportunities for improved clinical care through genomics-enabled medicine.

## Acknowledgements

This research has been conducted using the UK Biobank Resource under project number 24268. We appreciate the benevolence of UK Biobank volunteers.

